# A critical re-evaluation of multilocus sequence typing (MLST) efforts in *Wolbachia*

**DOI:** 10.1101/133710

**Authors:** Christoph Bleidorn, Michael Gerth

## Abstract

*Wolbachia* (Alphaproteobacteria, Rickettsiales) is the most common, and arguably one of the most important inherited symbionts. Molecular differentiation of *Wolbachia* strains is routinely performed with a set of five multilocus sequence typing (MLST) markers. However, since its inception in 2006, the performance of MLST in *Wolbachia* strain typing has not been assessed objectively. Here, we evaluate the properties of *Wolbachia* MLST markers and compare it to 252 other single copy loci present in the genome of most *Wolbachia* strains. Specifically, we investigated how well MLST performs at strain differentiation, at reflecting genetic diversity of strains, and as phylogenetic marker. We find that MLST loci are outperformed by other loci at all tasks they are currently employed for, and thus that they do not reflect the properties of a *Wolbachia* strain very well. We argue that whole genome typing approaches should be used for *Wolbachia* typing in the future. Alternatively, if few-loci-approaches are necessary, we provide a characterization of 252 single copy loci for a number a criteria, which may assist in designing specific typing systems or phylogenetic studies.

## Introduction

*Wolbachia* is a genus of maternally inherited intracellular Alphaproteobacteria that is found in arthropod and nematode hosts (Werren et al. 2008). Meta-analyses suggest that between 40% and 52% of all terrestrial arthropods are infected, making these bacteria the most common animal endosymbiont on earth (Zug & Hammerstein 2012; Weinert et al. 2015). Host specificity and type of symbiosis differs between major lineages of *Wolbachia*, which are currently classified into 16 supergroups named with capital letters from A–F and H–Q, consecutively in the order of their description (Glowska et al. 2015; Gerth 2016). Supergroups A and B are found in arthropods, representing the vast majority of described *Wolbachia* lineages. Many different types of symbioses, including reproductive parasitism, facultative mutualism, and obligate mutualism have been found for these lineages (Zug & Hammerstein 2015). In contrast, supergroups C and D are restricted to filarial nematodes, with which they share a close relationship that can be described as obligate mutualism (Makepeace & Gill 2016). Supergroup F has been found in both nematodes and arthropods and all other supergroups are rather rare, limited to a single or few hosts (Gerth et al. 2014).

Several host manipulations have been described for *Wolbachia,* and it is thought that those accelerate their spread in host populations, such as male-killing, feminization, induction of parthogenesis, and cytoplasmic incompatibility (Werren et al. 2008). These manipulations are considered to have a predominantly negative effect on their hosts. However, several positive aspects for hosts have been reported as well. These include provision of the host with amino acids or vitamins, or protection against viruses (Hedges et al. 2008; Teixeira et al. 2008; Zug & Hammerstein 2015). It appears likely that positive fitness effects drive the establishment of novel *Wolbachia* infections in host populations (Fenton et al. 2011; Kriesner et al. 2013). Recently, field studies demonstrated that mosquito populations can be artificially infected with fast spreading *Wolbachia* lineages which confer virus resistance to their hosts, thereby suppressing the transmission of the human pathogen Dengue (Hoffmann et al. 2011). However, not all strains of *Wolbachia* are able to confer virus resistance or to manipulate their host’s reproduction (Makepeace & Gill 2016).

The growing interest in the peculiar biology of *Wolbachia*, and its almost universal distribution among arthropods have necessitated means to differentiate strains by using molecular methods. Initially, genetic characterization of *Wolbachia* diversity was based on the 16S rRNA gene (O’Neill et al. 1992) or the more variable *wsp* gene (Zhou et al. 1998). However, in 2006, a multilocus sequence typing (MLST) system was established, and this subsequently became a standard in the community of *Wolbachia* researchers (Baldo et al. 2006).

The MLST approach was developed to provide a reproducible and portable method for the molecular characterization of bacterial pathogens. Originally designed to monitor local and global *Neisseria meningitides* outbreaks (Maiden et al. 1998), MLST schemes have since been published for many other bacterial species (Maiden 2006). For strain typing, five to ten loci (usually conserved housekeeping genes) from different regions of the genome are sequenced and each unique allele is assigned a unique number. Thus, a universal nomenclature based on a code of numbers referring to the sequenced loci is assembled. MLST genes are selected under the assumption that they underlie purifying selection, resulting in sequence variation that is mostly neutral. In the absence of recombination, substitutions should accumulate approximately linearly with time (Francisco et al. 2009) and therefore, genetic distances between strains at MLST loci would be proportional to their divergence time. MLST data are usually provided in a curated form in a freely accessible database (Jolley et al. 2004). Based on MLST profiles, relationships between (or diversity of) typed strains can either be analysed using the designated numbers from coding the alleles (i.e., MLST profiles), or by analysing the allelic nucleotide sequence data directly.

For *Wolbachia* MLST, fragments of five housekeeping genes (*gatB*, *coxA*, *hcpA*, *fbpA*, and *ftsZ*) are sequenced, and primers that amplify these loci across the major *Wolbachia* supergroups in arthropods are available (Baldo et al. 2006). According to the high number of citations for the original publication (Baldo et al. 2006, 343 citations in ISI Web of Science accessed August 17^th^, 2017), the approach is well-established and frequently used in the community of *Wolbachia* researchers. Since its original description more than 10 years ago, 2355 sequences and 472 unique MLST profiles have been added to the database (https://pubmlst.org/wolbachia/, accessed August 17^th^, 2017). When MLST was conceived, only two *Wolbachia* strains were represented by a fully annotated genome, and therefore, it was not possible to test how well MLST reflects the true *Wolbachia* strain diversity. Now, with a plethora of strains characterized by MLST, and several complete or draft genomic sequences of *Wolbachia* strains available (>30 strains in public repositories), the efficiency and performance of *Wolbachia* MLST can be evaluated objectively.

In this article, we aim to do so by first identifying the most common tasks *Wolbachia* MLST has been employed for by the research community. Using whole-genome as well as MLST data, we next assess how well MLST performs in these tasks in comparison to other single copy loci. We will argue that there is not a single locus or a single set of loci that performs well in all questions that are commonly addressed by *Wolbachia* researchers. Although the MLST scheme is convenient in that it provides a readily employable set of molecular markers, its information content is critically dependent on the research objective and the set of strains analysed. We therefore advocate that molecular markers for *Wolbachia* should be chosen very carefully for each particular research question, ideally based on whole-genome information.

### Usage of *Wolbachia* MLST in theory and research praxis

Originally, MLST was aimed to provide “a reliable system for typing and quantifying strain diversity” that allows “tracing the movement of *Wolbachia* globally and within insect communities and for associating *Wolbachia* strains with geographic regions, host features (e.g., ecology and phylogeny), and phenotypic effects on hosts” (Baldo et al. 2006). In other words, ideally each *Wolbachia* strain in the MLST database would not only be represented by a MLST profile, but also be linked with taxonomic information about its host, geographic origin, and phenotypic effects. This would then enable comparative analyses. However, out of 1828 strains (“isolates”) currently listed in the MLST database, only 603 (~34%) are associated with host taxonomy on the level of host order, and even fewer are associated with a host species (542, ~30%). Similarly, only 577 isolates (~31%) have geographic information and a phenotype is only known from 92 strains (~5%). Thus, the majority of *Wolbachia* strains in the database are defined by their MLST profiles alone, which further are in most cases incomplete (~60% of strains lack one or more alleles). Although this likely impedes comparative analyses, the lack of metadata associated with *Wolbachia* MLST isolates is not a problem for strain definition as such. However, if MLST is the only definition for a *Wolbachia* strain, it is crucial to understand how appropriate this definition is and to ascertain that the MLST profile is not isolated from the biological properties of the typed strains.

In current practise, it is generally assumed that MLST markers are a good approximation of genome-wide characteristics of *Wolbachia* strains. As such, they have been used to describe and analyse the *Wolbachia* diversity, phylogeny, or phylogeography of particular host taxa (Russell et al. 2009; Watanabe et al. 2012; Schuler et al. 2013; Zhang et al. 2013a; Sontowski et al. 2015), taxa from a particular ecological background/community (Stahlhut et al. 2010; Zhang et al. 2013b), and to explore horizontal movements of *Wolbachia* strains (Baldo et al. 2008; Gerth et al. 2013; Ahmed et al. 2016). All of these research questions entail a number of implicit assumptions about the performance of *Wolbachia* MLST. We will in the following examine three of these assumptions that we consider most important in this regard:

1) *Wolbachia* MLST can differentiate *Wolbachia* strains.

2) Genetic divergence at *Wolbachia* MLST genes corresponds to genome–wide divergence levels

3) *Wolbachia* MLST gene phylogeny reflects the phylogeny of the core genome.

### Differentiating *Wolbachia* strains with MLST markers

One common task for MLST in *Wolbachia* research is the discrimination (or “quantification” as in Baldo et al. 2006) of *Wolbachia* strains, i.e., to answer if two (or any other number) of strains are genetically different. For this task, the level of resolution depends on the number and type of genes used, length of the sequences and the genetic diversity of chosen loci (Cooper & Feil 2004). The limits of MLST schemes were pointed out for genetically monomorphic bacteria such as *Mycobacterium tuberculosis* or *Bacillus anthracis* (Achtman 2008; Achtman 2012). *Wolbachia* MLST diversity within supergroups is far from being monomorphic, as evident from the large number of available profiles in the database (see above). However, the actual evolutionary pace of *Wolbachia* genes and genomes was an open question. Recently, based on a time-calibrated phylogenomic analyses it was hypothesized that *Wolbachia* lineages are much older than previously assumed – and therefore that genetic change due to substitutions or recombination accumulate slower than expected (Gerth & Bleidorn 2016). In accordance with this estimate, it was repeatedly reported that *Wolbachia* MLST is not suited to discriminate between closely related strains (Ishmael et al. 2009; Atyame et al. 2011; Riegler et al. 2012; Siozios et al. 2013a; Conner et al. 2017). This does not come as a surprise, as per definition, MLST genes are of conserved nature, and thus slowly evolving. They are therefore inherently unsuited to trace very recent evolutionary events.

Comparing the ability of MLST loci to differentiate *Wolbachia* strains with that of 252 other single copy loci employed in a recent phylogenomic study of *Wolbachia* evolution (Gerth & Bleidorn 2016) shows that MLST loci are not ideal for this task (Supplementary Table 1). MLST loci are able to differentiate 42–63% of the 19 analysed *Wolbachia* genomes, whereas other conserved single copy loci may differentiate up to 84% of the strains (16/19). *Wsp* is more variable than MLST markers (13/19 strains differentiated), but is outperformed by a number of single copy loci (Table S1). Strikingly, none of the 252 loci that were originally selected as phylogenetic markers can discriminate between all strains. This is because they were chosen to be present in a single copy in all of the analysed *Wolbachia* genomes (Gerth & Bleidorn 2016) and thus also represent mostly conserved housekeeping loci (Supplementary Table 1). In summary, conserved single copy genes are generally unsuited markers to differentiate for closely related *Wolbachia* genomes, and among those, MLST and *wsp* loci do not perform particularly well.

Therefore, when designing an experiment with the main or foremost goal of differentiating *Wolbachia* strains, one should employ fast evolving markers such as ankyrin repeats, insertion sequences, or other mobile elements that have been shown to be the fastest evolving genomic features of *Wolbachia* (Wu et al. 2004; Tanaka et al. 2009; Newton et al. 2016). As these will likely be very different between distantly related strains (Cerveau et al. 2011), a universal set of markers suitable across the breadth of *Wolbachia* diversity does not exist. As a consequence, in many cases it will be inevitable to identify suitable markers for *Wolbachia* differentiation through comparative genomics of a representative sample of the strains to be investigated.

Furthermore, we advocate to adjust not only the type, but also the number of loci employed for strain differentiation. Random sampling MLST profiles from the known diversity of *Wolbachia* MLST profiles illustrates that in many cases, two or three MLST loci provide similar resolution to all five MLST genes (Fig. 1). For example, when analysing 20 *Wolbachia* strains and using only the two most variable MLST genes *hcpA* and *gatB*, one would on average be able to differentiate at least 19 of these strains. For 40 strains, three loci provide a similar resolution (Fig. 1). Although this comes with the caveat that not all systems will show the same *Wolbachia* MLST profile frequencies as the MLST database, it demonstrates that careful adjustment of loci to the study system can save time and money. Instead of typing all *Wolbachia* samples with five MLST loci, we therefore recommend to maximise the number of detectable *Wolbachia* strains by first typing with the fastest evolving marker available (ideally, this would have been identified *a prior*i through comparative genomics), and then continue with additional markers as the number of samples increases.

**Fig. 1.**
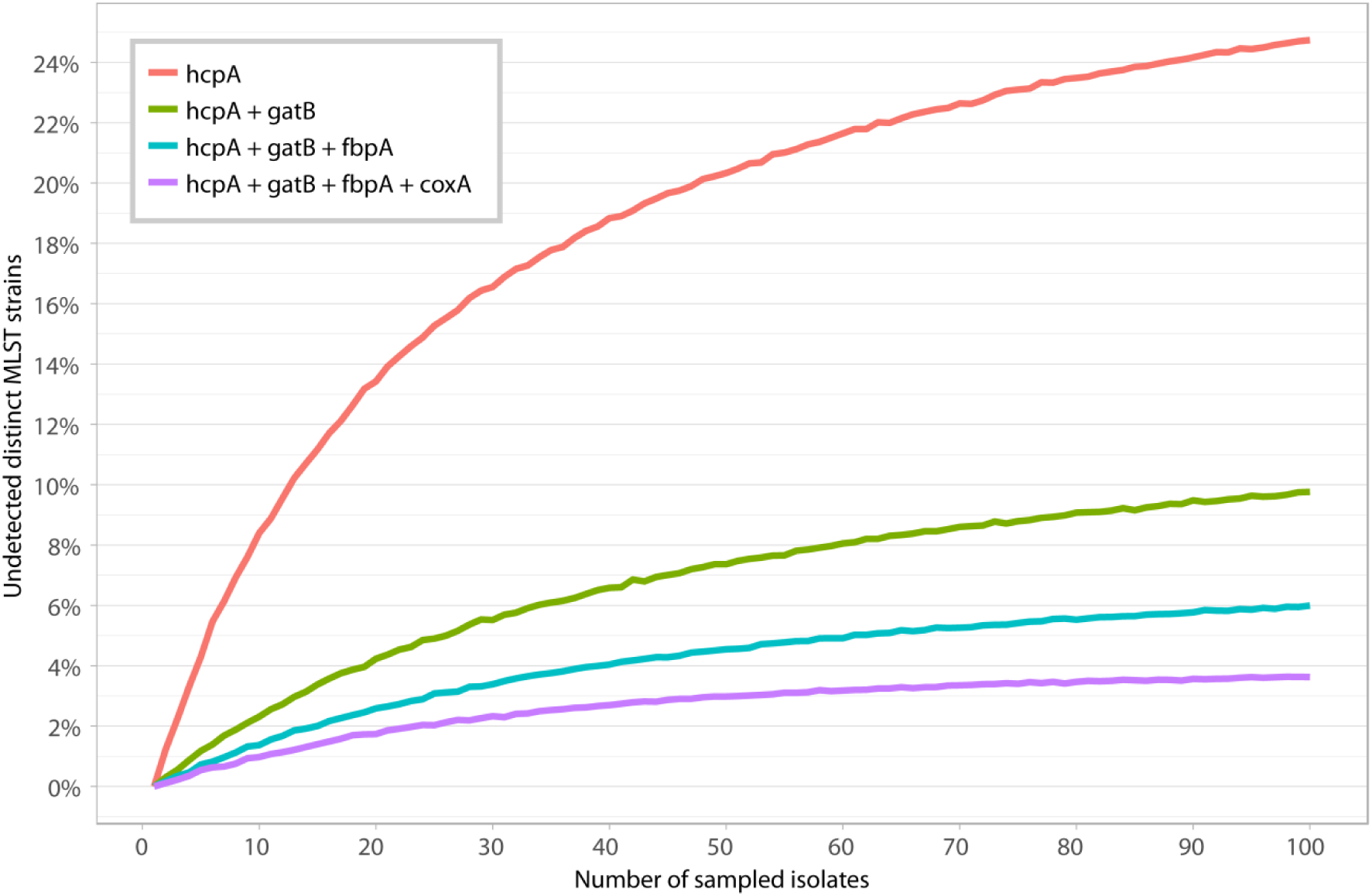
Ability of MLST markers in differentiating *Wolbachia* strains. Average proportion of undetected distinct *Wolbachia* MLST profiles when using only one, two, three or four MLST genes is displayed in relation to the number of analysed strains. Figure is based on all complete *Wolbachia* MLST profiles (740 in total, 472 of which are unique) currently available from the pubMLST database (https://pubmlst.org/wolbachia/, last accessed August 17^th^, 2017). See methods for details.

### Assessing genetic differentiation of *Wolbachia* strains with MLST genes

In addition to differentiating strains, a strain typing system should also be able to characterize the genetic diversity of a set of strains to be analysed. For this to be as accurate as possible, the molecular divergence of investigated strains at their MLST loci would have to be identical or very similar to genome wide divergence rates, or correlate with genome wide rates very well. If the assumptions underlying the choice of MLST loci (mostly neutral selection, see above) are correct, one might expect these two characteristics to be met. However, an analysis of MLST vs. core genome divergence rates shows that this is not true for the currently employed *Wolbachia* MLST loci (Fig. 2). As expected from the previous observations (see above) core genome divergence of lower than ~0.2% cannot be detected with any of the MLST loci (Fig. 2). For *ftsZ*, even strains that are genetically divergent by more than 1% may appear identical.

**Fig. 2.**
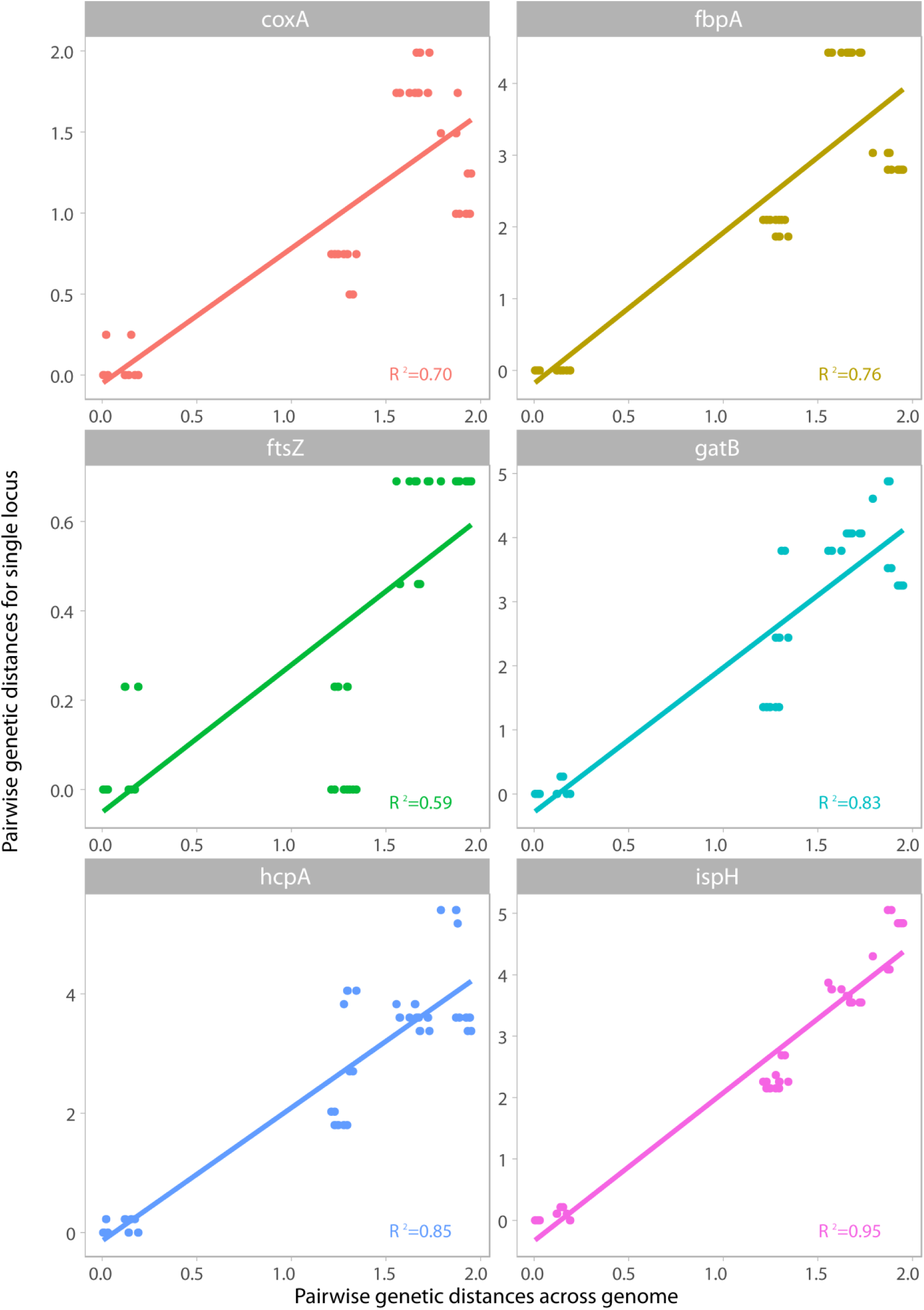
Correlation of genetic distances of *Wolbachia* strains at MLST loci to genome-wide distances. Each data point corresponds to a single pair of *Wolbachia* strains, and shows the divergence between these two strains at MLST loci (y-axis) and genome-wide distance (x-axis, mean distance from 252 single copy orthologs). Panels correspond to one of the 5 MLST loci and *ispH* (encoding 4-hydroxy-3-methylbut-2-enyl diphosphate reductase) for comparison. Linear regression models were fitted using the R statistical environment (R Core Team 2015). All distances are displayed as raw genetic distances in percent. Please note that all pairwise distances are from supergroup A strains only, as including supergroup B strains would lead to skewed distributions (small distances within supergroups and large distances between supergoups) and therefore biased correlation estimates. All R^2^ values for all analysed loci and both supergroups can be found in Supplementary Table 1. Correlations of divergence at *wsp* vs core genome loci can be found in Supplementary Fig. 1.

Furthermore, a number of strains that are diverged by 1.5–2% appear similarly divergent at their *ftsZ* and *fbpA* loci (Fig. 2). This may indicate nucleotide substitution saturation, which would impede genetic comparison of distantly related *Wolbachia* strains with MLST.

Further to these patterns, out of the five MLST loci, only *coxA* shows genetic divergence rates similar to those obtained from whole genome information, whereas those of *ftsZ* are lower and the ones from *hcpA*, *fbpA* and *gatB* are higher (Fig. 2).

Finally, none of the divergence rates estimated from the 5 MLST loci correlate very well with genome wide rates (R^2^ values of regression in linear model 0.59–0.85, Fig. 2), which contrasts with loci that show a very good correlation in this respect (e.g., *ispH*, Fig. 2). For *wsp*, the relation of genetic distances to core genome distances can be described as random (Supplementary Fig. 1). In summary, the MLST loci are not a good approximation for genome wide divergence rates of *Wolbachia* strains, and other loci may be more appropriate (Fig. 2, Supplementary Table 1). This also means that genetic divergence ratios obtained from MLST loci should be interpreted cautiously and other loci should be explored as alternative. However, as the performance at this task differs even for a single locus between supergroups (Supplementary Table 1), comparative genomics will again in many cases be the only option to reliably determine divergence rates between a sample *Wolbachia* strains.

### Phylogenetic analyses of *Wolbachia* strains using MLST

Judging from the abstracts and keywords of all articles citing the original *Wolbachia* MLST publication, questions that are commonly addressed with *Wolbachia* MLST are phylogeny & phylogeography (102 articles with corresponding terms), and horizontal transmission of strains (58 articles). Since the determination of horizontal transmission with molecular methods also requires phylogenetic approaches, one can summarize that phylogenies are one major field of application for *Wolbachia* MLST.

This is despite the authors’ original assessment that MLST loci are not necessarily good phylogenetic markers (“caution in interpretation of phylogenetic relationships is necessary”, Baldo et al. 2006), and despite the fact that an assessment of its performance as phylogenetic marker as lacking.

The level of resolution across time for a given gene in a phylogenetic analysis can be estimated by its phylogenetic informativeness (PI), which measures the relative ratio of phylogenetic signal to noise across time (Townsend 2007). Analysing the PI profiles of all MLST genes for a set of *Wolbachia* strains covering supergroup A and B reveals that all of them show the highest phylogenetic resolution on the supergroup level (Fig. 3). According to Gerth & Bleidorn (2016), the supergroups A and B have diverged more than 200 million years ago. MLST genes however provide only little phylogenetic information for strains that diverged much more recently (Fig. 3). As *Wolbachia* likely moves between hosts at a fast rate (Gerth et al. 2013; Bailly-Bechet et al. 2017), the MLST approach is not suited to infer phylogenetic relationships of closely related strains, to detect recent horizontal transmissions or to assess ecological timescales of *Wolbachia* movements between populations. However, a number of *Wolbachia* genes–including the highly recombining *wsp*-evolve considerably faster than MLST loci (as measured by genetic divergence or number of variable alignment sites) and also provide phylogenetic information on very shallow phylogenetic levels (Fig. 3, Supplementary Table 1, Supplementary Fig. 2). These loci might be good candidates for resolving very recent evolutionary events.

**Fig. 3.**
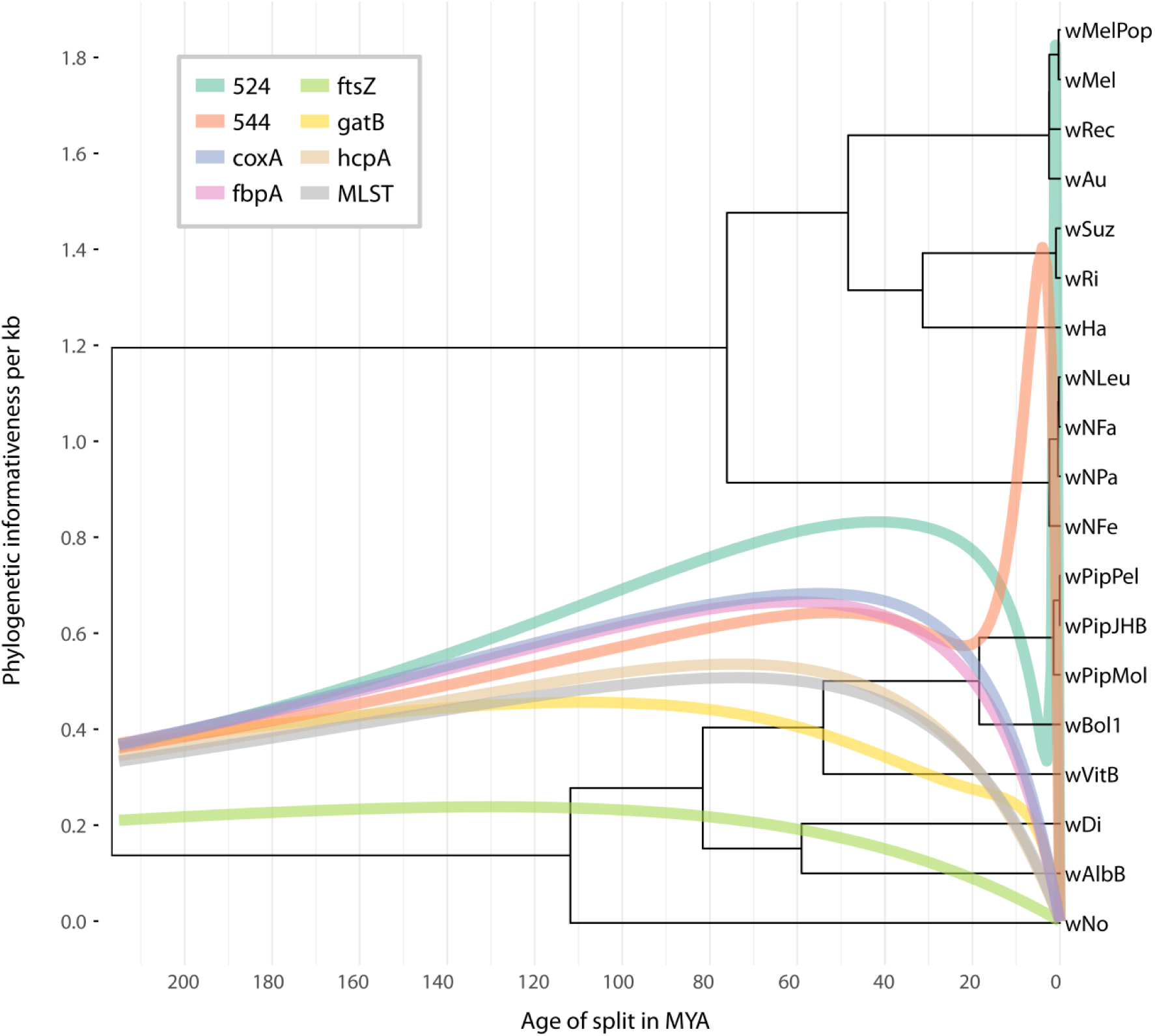
Phylogenetic informativeness (PI) of five MLST-gene alignments from supergroup A and B *Wolbachia* strains. For comparison, two loci displaying relatively high PI for recent evolutionary events are also shown. Ultrametric tree (based on 252 loci available under https://github.com/gerthmicha/wolbachia-mlst/tree/master/alignments) was taken from Gerth & Bleidorn (2016). See Supplementary Figure 1 for PI profiles for of all analysed loci.

Moreover, as already mentioned in the original MLST publication (Baldo *et al.*, 2006), all MLST loci except for *ftsZ* show some level of intragenic recombination. Indeed, using all alleles present in the *Wolbachia* database today, signals of recombination can be detected for all five markers using the PHI test (Bruen et al. 2006). The presence of horizontal genetic exchange makes the interpretation of phylogenetic analyses of MLST genes challenging, as the resulting tree may not reflect the evolutionary relationships of the analysed strains (Holmes et al. 1999; Jiggins et al. 2001). When comparing the phylogenetic reconstruction for the dataset in Fig. 3 using the five concatenated MLST-fragments with the original analysis based on 252 orthologs, several differences in the topology are apparent (Fig. 4A, B). Seven internal nodes are reconstructed differently in the MLST-based analysis (Fig. 4B) and the branch length differences between analyses, especially within supergroups are striking. These differences are likely due to the misleading signal from recombination events.

**Fig. 4.**
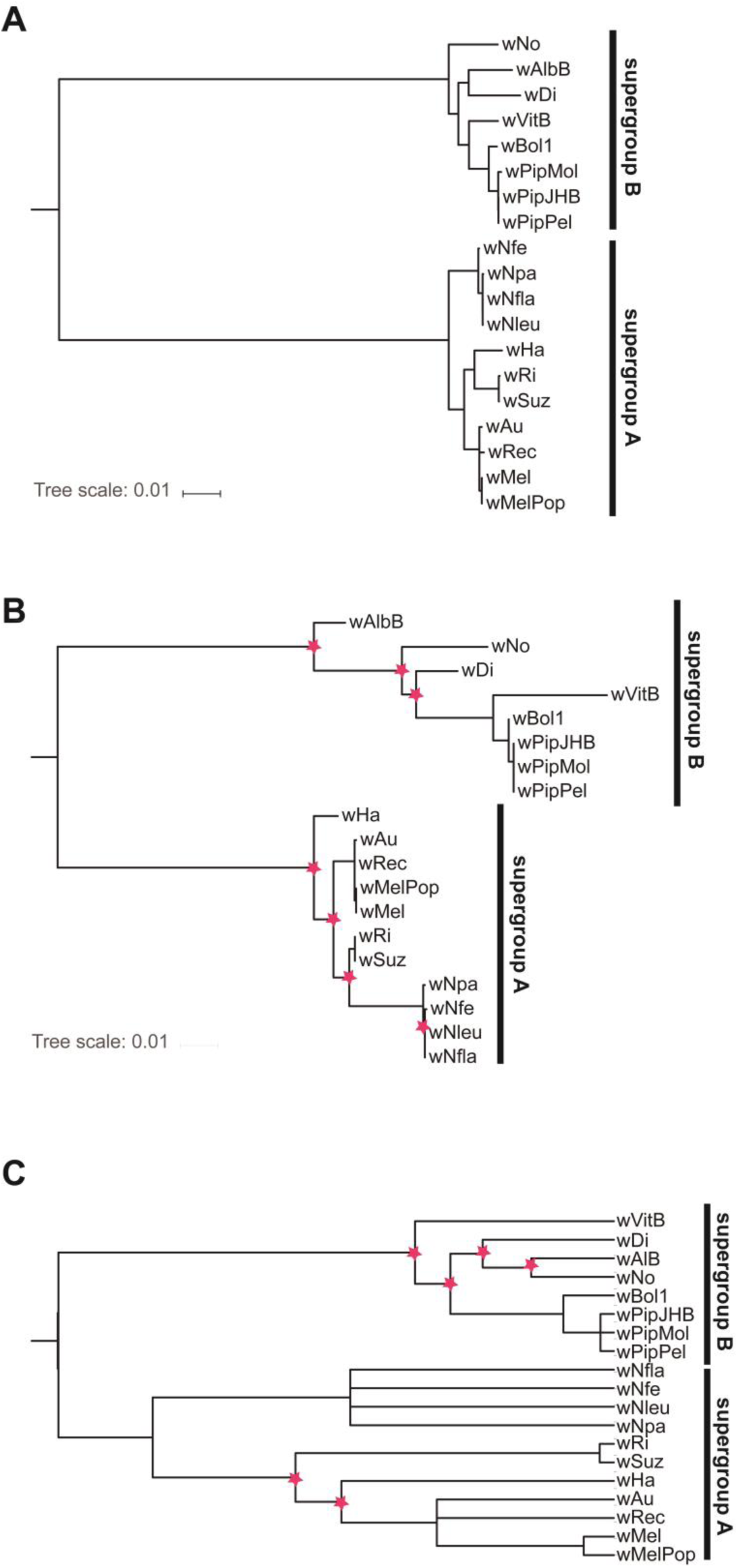
Phylogenetic reconstruction of supergroup A and B strains as selected by Gerth & Bleidorn (2016). A) Maximum likelihood analysis based on optimal partitions and models as selected by IQ-TREE (Nguyen et al. 2015) for a dataset containing nucleotide data of 252 non-recombining orthologs. B) Maximum likelihood reconstruction of five MLST gene fragments of the same taxa using optimal partitions and models as selected by IQ-TREE. Seven conflicting nodes (red asterisks) compared to the phylogenomic analyses are highlighted. C) ClonalFrame (Didelot & Falush 2007) analysis of the MLST dataset, with six conflicting nodes highlighted (red asterisks).

ClonalFrame is a Bayesian phylogenetic framework that was developed especially for MLST datasets and capable of inferring relationships despite the presence of recombination (Didelot & Falush 2007). Nevertheless, analysing our dataset with ClonalFrame led to a similarly high number of conflicting nodes (six) in comparison to the phylogenomic dataset, and multiple polytomies (i.e., unresolved nodes, Fig 4C). This shows that the usage of recombination-aware phylogenetic methods cannot circumvent the inherent problems of *Wolbachia* MLST genes as phylogenetic markers. As some level of conflict exists between the trees recovered from most single gene loci and the one from the supermatrix (Supplementary Fig. 2), whole genome based phylogenies are desirable to minimize biases.

Homologous recombination is widespread among Bacteria (Didelot & Maiden 2010). One way to circumvent problems in phylogenetics arising from recombination is to estimate relationships between strains based on allele designations. A simple method for this is to cluster strains based on their similarity, which can be visualized as a dendrogram. However, strain similarity does not necessarily reflect common ancestry. A popular and more sophisticated method to analyse allele-based strain data is eBURST (Feil et al. 2004). This software incorporates a model of bacterial evolution in which strains that are increasing in frequency diversify, thereby forming clusters of similar genotypes. For MLST data, so-called clonal complexes are defined as groups that share a predefined number of alleles (e.g., three of five allele designations are identical) with at least one other strain type. After searching for these clonal complexes, the likely founding strain type is inferred, as are evolutionary relationships within this clonal complex. Simulation studies have shown that when recombination is absent or present in low to moderate levels, the inferred relationships of clonal complexes are very similar to the (known) true ancestry (Turner et al. 2007). However, increasing rates of the frequency of recombination to mutations led to a strong decrease in the reliability of eBURST analyses. In *Wolbachia,* the overall ratio of recombination to mutation events to explain the generation of a substitution is ranging from 2.3 to 8.2, depending on the analysed genome (Ellegaard et al. 2013). Therefore, the high recombination rates in *Wolbachia* genomes make allele-based analyses unreliable.

In addition to problems with recombination, there are also theoretical arguments against using ‘eBURST’-like clustering algorithms with *Wolbachia* MLST profiles. Because the only criterion for assigning a novel allele number is at least one nucleotide difference compared to all described alleles, any number of substitutions in one allele is weighted equally. For example, 10 different *Wolbachia* strains may be differentiated by only 9 nucleotide differences in total, or by 50, and could potentially be characterised by identical sets of MLST profiles. This makes comparing these profiles across systems challenging. When sampling is dense and therefore the majority of the allele diversity is known, this will likely not be problematic. However, this is rarely ever the case for *Wolbachia.* Given the large number of infected species, it is essentially impossible to know the true diversity of *Wolbachia* in any ecosystem. Furthermore, because horizontal transmissions are common (Baldo et al. 2008; Zug et al. 2012; Gerth et al. 2013; Ahmed et al. 2016), and exact pathways of these transmissions are still discussed (Huigens et al. 2004; Le Clec’h et al. 2013; Li et al. 2016), it does not make sense to define “founding” and “descending” *Wolbachia* genotypes in most cases.

### Alternatives to MLST

MLST was developed as a replacement for an earlier strain typing approach called multilocus enzyme electrophoresis (MLEE), which measured genetic variation by the resolution of electrophoretic variants (electromorphs) of metabolic enzymes (Maiden 2006). One problem of this method was that experiments were difficult to reproduce across labs. With the availability of affordable and faster Sanger sequencers it was possible to directly use sequence data instead of electromorphs. Nowadays, a wide array of different high-throughput sequencing techniques is available (Bleidorn 2015; Goodwin et al. 2016). Due to their small size, sequencing of complete bacterial genomes is affordable and routinely carried out using benchtop sequencers in laboratories with standard equipment (Loman & Pallen 2015). Consequently, strain typing methods based on whole genome data were proposed, e.g., rMLST, in which a set of 53 ribosomal proteins is used (Jolley et al. 2012). Ribosomal proteins are universally found in bacterial genomes, show a wide distribution across genomes and are expected to underlie stabilizing selection, similar to the above mentioned MLST genes. In the case of *Wolbachia*, ribosomal proteins have already been used successfully for phylogenomic analyses (Nikoh et al. 2014). Other typing methods simply employ all available genes, i.e., whole genome sequence-typing (WGST) (Pérez-Losada et al. 2013) or core genome MLST (cgMLST) (De Been et al. 2015).

Although *Wolbachia* harbour small genomes (1 to 1.5mbp in size) (Makepeace & Gill 2016), sequencing and assembly is more difficult than for many other Bacteria. As strictly intracellular endosymbionts, *Wolbachia* cannot be cultured axenically, and although maintaining them in cell cultures is possible (Dobson et al. 2002), it is very laborious and often not practical. Thus, in many cases a metagenomic sequencing approach is used, targeting both host and *Wolbachia* DNA. *Wolbachia* sequence data can then be retrieved using BLAST-searches and read mapping (Gerth et al. 2014). However, in this case a high sequencing depth per genome is needed, as typically only a small proportion of the reads will be of *Wolbachia* origin. For more efficient sequencing of *Wolbachia* genomes, target enrichment protocols (Lemmon & Lemmon 2012) have been established (Geniez et al. 2012; Dunning-Hotopp et al. 2017), although these are not yet broadly applied.

Another problem in *Wolbachia* genome sequencing is the high density of mobile genetic elements with repetitive sequence motives (Wu et al. 2004), which may lead to very fragmented assemblies. However, for analyses focussing on sequence data of selected loci and not on synteny, incompletely assembled *Wolbachia* draft genomes are sufficient. Working with complete (or draft) genomes has the advantage that comparative analyses can be used to retrieve large sets of orthologous and recombination-free loci (Comandatore et al. 2013). These datasets allow to circumvent almost all problems with MLST outlined in this article, and further enable the identification hypervariable regions such as tandem repeat markers (Riegler et al. 2012) or ankyrin repeat domains (Siozios et al. 2013b).

Although whole genome approaches are the arguably the best way to address *Wolbachia* strain differentiation, diversity estimates, and phylogeny, they may in some cases be too cost-or time intensive, and there will be questions that must be addressed with a small number of genetic marker loci. In this case we here provide a characterization of 252 conserved single copy genes by a number of criteria, each of which may be important in strain typing, depending on the question to be addressed (Supplementary Table 1). We point out that for none of these criteria, the MLST loci perform particularly well, and we therefore strongly suggest to chose marker loci based on the experimental design rather than on the convenient availability

### Summary & conclusion

MLST analyses are widely used in the community of *Wolbachia* researchers and a large database for comparative studies is available. This database and the availability of PCR protocols for most *Wolbachia* strains represent a convenient and valuable resource. However, for most tasks routinely employed for, *Wolbachia* MLST markers are unsuited. They are too conserved to allow reliable and fine-scaled strain differentiation, they do not reflect genome wide divergence rates well, and they are poor phylogenetic markers at shallow or deep divergence levels. Further, they are outcompeted at all of these tasks by other loci. These properties make the definition of a strain in the genus *Wolbachia* per MLST very problematic and we recommend that this practice is discontinued. Instead, we advise to tailor adequate marker loci as required for the investigated strains. Naturally, these will differ between study systems and research questions, but we think that the shortcomings of MLST loci outweigh their benefit of universality. Generally, we hope that the *Wolbachia* community will embrace whole genome typing methods, which are already standardly employed in clinical microbiology. However, efficient novel *Wolbachia* genome sequencing (or enrichment) protocols are needed for this to succeed.

## Methods

### Data acquisition

Most MLST sequences, isolates and profiles described and analysed in this paper were downloaded from the *Wolbachia* PubMLST database (Jolley et al. 2004; Baldo et al. 2006; https://pubmlst.org/wolbachia/, last accessed 17th of August 2017). For comparative analysis of 19 supergroup A and B *Wolbachia* strains, the corresponding MLST gene sequences were recovered via blastn (Camacho et al. 2009) searches against coding nucleotide sequences of the 19 *Wolbachia* strains, using MLST sequences from the online database as a query. The hits were trimmed manually to conform to the length of *Wolbachia* MLST alleles. In addition, 252 loci from complete or draft *Wolbachia* genomes were acquired as described in Gerth & Bleidorn (2016). Briefly, the 252 loci were single copy genes present in all of the 19 investigated *Wolbachia* strains that did not show evidence for recombination. Orthology was assessed with OrthoFinder version 0.2.8 (Emms & Kelly 2015), and alignment was performed based on codons using Mafft version 7.215 (Katoh & Standley 2013). In the following, the performance of *Wolbachia* MLST loci was compared to that of the 252 loci with regard to their ability to differentiate strains, to approximate genome-wide divergence, and to reflect core genome phylogeny.

For the sake of completeness, these comparisons also included *wsp* (*Wolbachia* surface protein). Although not very commonly in use today, it was suggested as additional marker in *Wolbachia* typing schemes (Baldo et al. 2006) and was the standard molecular marker for *Wolbachia* before the development of MLST (Zhou et al. 1998).

However, it was repeatedly pointed out that *wsp* is not a suitable marker for molecular typing of *Wolbachia* strains (Paraskevopoulos et al. 2006; Baldo & Werren 2007).

### Strain differentiation

Strain differentiation ability was assessed for all investigated loci by the proportion of distinct alleles in all alleles. This was calculated using the function 'haplotype' of the R package pegas (Paradis 2010; R Core Team 2015). As additional measures of strains differentiation, we calculated average pairwise genetic distances and the number of variable alignment sites using the functions 'dist.dna' and 'seg.sites' of the R package APE (Paradis et al. 2004), respectively. All measures can be found in Supplementary Table 1.

To determine the resolution of the single, two, three or four most variable MLST loci in comparison to all five loci, we randomly sampled MLST profiles from the known diversity of MLST strains in the pubMLST database (at the time of the analysis, 740 complete MLST profiles, 472 of which were unique). Random sampling was performed for datasets of 1–100 samples, and repeated 10,000 times in all cases. The number of distinct isolates among the samples based on a single, two, three or four MLST loci was counted and compared to the number of distinct isolated based on complete MLST profiles.

### Divergence rates

For all investigated loci, we aimed to assess how well genetic distances of a single locus reflect the genetic distances of the core genome. To this end, we calculated all possible pairwise raw genetic distances (55 pairwise distances for 19 strains analysed) for each MLST locus, *wsp*, and for the concatenated 252 loci (as approximation of the core genome) as described above. Next, the correlation of the distances from each single locus with the core genome was determined by fitting a linear model within the R statistical framework. All R^2^ values for these models can be found in Supplementary Table 1. Due to the nature of the dataset, there is a bimodal distribution of distances: large distances between supergroups, and small distances within supergroups. Using this biased dataset, all correlation measures for all loci were very high. Therefore, we decided it would be more appropriate to perform this analysis separately for each supergroup.

### Phylogenetic analyses

Phylogenetic analyses of 19 *Wolbachia* strains was performed for a dataset of five concatenated MLST genes, one dataset of 252 concatenated single copy orthologs and for each of the 258 investigated loci (5 MLST genes, 252 core genome loci, *wsp*) separately. For all analyses, a maximum likelihood tree search was performed with IQ-TREE version version 1.5.4 (Nguyen et al. 2015) using the implemented optimal model search and, for multi-gene analyses, optimal partition selection algorithms (Lanfear et al. 2012; Chernomor et al. 2016; Kalyaanamoorthy et al. 2017). The MLST dataset was further analysed with ClonalFrame version 1.2 (Didelot & Falush 2007), using four independent runs with 1,000,000 generations each and a burnin of 50% for all runs. Convergence of runs and stability of sampled parameters was verified by plotting likelihood values and other parameters in R. All runs converged on identical topologies. Congruence and conflict between single gene analyses and core genome analysis was also assessed by calculating normalized Robinson-Foulds distances (Robinson & Foulds 1981) with RAxML version 8.2.1 (Stamatakis 2014) between single gene trees and the tree that best represented core genome phylogeny. Additionally, we calculated the likelihood of each single gene topology with RAxML using the 252 loci dataset. Congruence was approximated by calculating the difference between core genome topology log likelihood and the likelihoods of each single gene analysis

Finally, phylogenetic informativeness (PI), i.e., the relative amount of phylogenetic signal to noise across time was estimated for all analysed loci using

TAPIR (Faircloth et al. 2012), an efficient implementation of Townsend’s phylogenetic informativeness (Townsend 2007), which makes use of the HyPhy software package (Pond & Muse 2005). To this end, an ultrametric tree of the analysed *Wolbachia* strains was taken from (Gerth & Bleidorn 2016).

## Funding

This work was supported by the Spanish Ministry of Science and Education (MEC) [RYC-2014-15615 to CB]; European Molecular Biology Organization [ALTF 48-2015 to MG]; and Marie-Curie Actions of the European Commission [LTFCOFUND2013, GA-2013-609409 to MG].

